# Spatiotemporal Network Dynamics Reveal Alzheimer’s Disease Progression

**DOI:** 10.64898/2025.12.09.692653

**Authors:** Theodore J. LaGrow, Vaibhavi Itkyal, Harrison Watters, Kyle M. Jensen, Ram Ballem, Wen-ju Pan, Armin Iraji, Vince D. Calhoun, Shella Keilholz

## Abstract

Alzheimer’s disease (AD) is characterized by progressive disruptions in large-scale brain networks that precede cognitive decline, yet conventional functional connectivity analyses often fail to detect disruptions in coordination among large-scale brain networks that may be critical for early detection. This study leverages quasi periodic patterns (QPPs) and complex principal component analysis (cPCA) to characterize spatiotemporal network alterations across longitudinally stable (normal cognitive, mild cognitive impairment, dementia of Alzheimer’s type) and transitioning (normal cognitive to mild cognitive impairment, mild cognitive impairment to dementia of Alzheimer’s type) cohorts from the Alzheimer’s Disease Neuroimaging Initiative using resting state fMRI. QPPs were used to derive recurrent spatiotemporal templates and network integrity measures at the intrinsic connectivity network level, while cPCA decomposed Hilbert transformed time series into complex valued patterns that capture amplitude and phase relationships. Nonparametric group comparisons revealed a structured trajectory in which limbic, subcortical, and higher cognition networks, including triple network components, are affected early, followed by progressive disruption in visual, cerebellar, sensorimotor, and additional triple network systems. Transitioning cohorts showed many of these alterations before formal diagnostic conversion, indicating that spatiotemporal signatures carry preclinical information. QPP based metrics were particularly sensitive to limbic and subcortical degradation, whereas cPCA emphasized changes in higher order, visual, and cerebellar patterns, revealing complementary aspects of the same underlying pathology. These findings extend prior QPP only work and highlight the utility of combining QPP and cPCA based measures as a dynamic, network-level biomarker framework for AD progression. with potential applications in early detection, characterizing disease trajectories, and treatment monitoring.

## I. INTRODUCTION

Alzheimer’s disease (AD) is a leading cause of dementia and a major global health burden, with tens of millions of individuals affected and projections indicating a substantial rise in prevalence and cost over the coming decades [1]. As life expectancy increases, so does the number of people at risk for cognitive decline, creating an urgent need for noninvasive tools that can detect and track AD related brain changes before severe impairment develops [1].

AD pathology accumulates long before overt cognitive symptoms emerge. Conventional static functional connectivity (FC) analyses in functional MRI (fMRI) summarize average coupling, thereby averaging out the rich temporal structure of ongoing brain activity. Transient, recurrent perturbations in large scale networks that may signal early neurodegenerative change can therefore be obscured [2]–[4]. Capturing these spatiotemporal patterns is a key step toward more sensitive functional biomarkers of AD.

Quasi periodic patterns (QPPs) offer one way to characterize such dynamics. QPPs are recurrent, phase coupled spatiotemporal patterns in resting state fMRI. Prior work shows that QPP expression is altered across several neuropsychiatric and neurodegenerative conditions, including transgenic rodent models of AD in which reduced QPP occurrence and disrupted spatial organization outperform conventional connectivity metrics for distinguishing disease from controls and correlate with amyloid pathology [5]. These findings suggest that QPPs may serve as dynamic markers of AD related network dysfunction.

Extending this work to humans, our previous analysis of resting state fMRI from the Alzheimer’s Disease Neuroimaging Initiative (ADNI) used QPP derived network integrity measures to track clinical stages spanning normal cognition (NC), mild cognitive impairment (MCI), and dementia of the Alzheimer type (DAT), as well as transitioning diagnoses [6]. That study demonstrated that QPP measures systematically tracked disease stage, supporting QPPs as a candidate functional biomarker for AD,

QPPs capture only one facet of the dynamics: a recurrent spatiotemporal template and its amplitude of expression over time. By adding complex principal component analysis (cPCA), we obtain a complementary, largely parameter free decomposition of the same large scale fluctuations, in which Hilbert transformed fMRI time series are represented as complex valued components whose spatial loadings and phase relationships explicitly encode traveling wave like patterns and cross network timing [7]. Unlike QPPs, which require specifying a template window length and correlation threshold, cPCA emphasizes the underlying variance structure and phase gradients of these modes, providing new insight into how amplitude and phase of disease related trajectories evolve across networks and clinical stages.

In the present study, we build directly on our prior QPP based analysis by leveraging QPPs and cPCA to investigate spatiotemporal network alterations across a spectrum of AD progression, including NC, MCI, and DAT, following the nomenclature of [8]. In addition to subjects with stable diagnoses, we examine individuals who transition between diagnostic categories, allowing us to probe early changes near clinical conversion. This extension incorporates cPCA derived QPP like patterns as a complementary representation of disease related dynamics and explicitly relates QPP and cPCA based metrics at the intrinsic connectivity network (ICN) level.

### This study addresses three questions

(1) Can QPP and cPCA based measures differentiate AD stages (NC, MCI, DAT) and detect preclinical changes in transiting cohorts, extending our prior QPP-only work [6]? (2) Which functional brain networks degrade earliest and most severely across the AD spectrum? (3) Do QPP and cPCA provide complementary rather than redundant information about large-scale network dynamics?

## II. METHODS

### A. Datasets

This study utilizes two groups from ADNI [9]. We categorized subjects into two primary groups based on disease progression and data availability. Group 1 consists of stable cohorts (sNC, sMCI, sDAT), defined as subjects who did not transition between disease stages during the ADNI study. Group 2 includes transitioning cohorts, specifically unstable normal cognition (uNC) and progressive mild cognitive impairment (pMCI) subjects, each with scans acquired before and after a diagnostic transition (Fig. 1B).

**Fig. 1:**
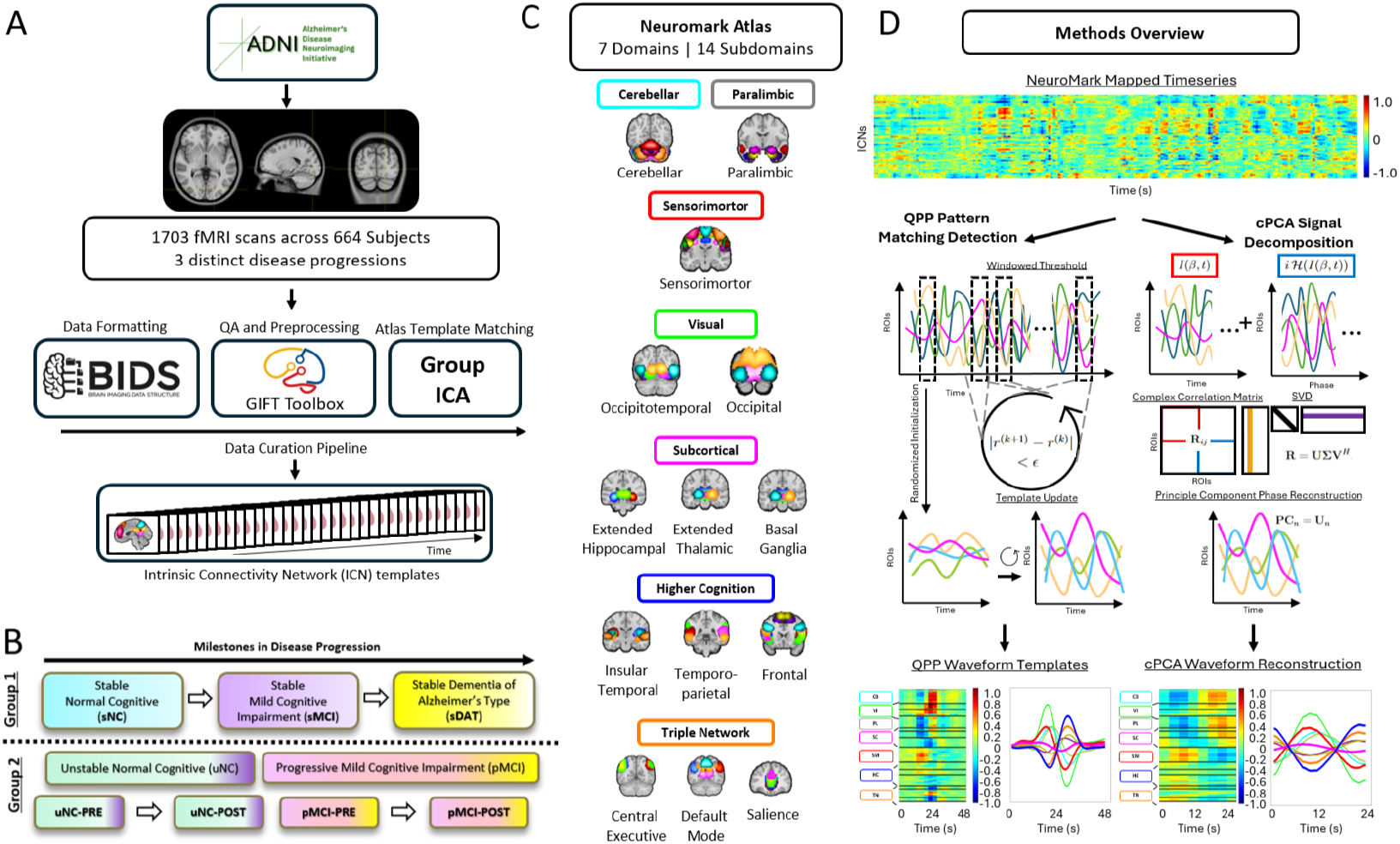
Overview of Spatiotemporal Projection. (A) Preprocessing pipeline for the ADNI [9] dataset into ICNs. (B) Experimental groups delineated as Group 1 (sNC, sMCI, sDAT) and Group 2 (uNC-PRE, uNC-POST, pMCI-PRE, pMCI-POST). (C) ICNs of the NeuroMark v2.2 Atlas [10], [11]. Each of the seven Domains are highlighted in color and the 14 Subdomains are listed below each coronal image depicting the 105 ICNs. The following are the 14 Subdomains: cerebellar (CB-CB), paralimbic (PL-PL), sensorimotor (SM-SM), visual-occipital (VI-OC), visual-occipitotemporal (VI-OT), subcortical-extended hippocampal (SC-EH), subcortical-extended thalamic (SC-ET), subcortical-basal ganglia (SC-BG), higher cognition-insular-temporal (HC-IT), higher cognition-temporoparietal (HC-TP), higher cognition-frontal (HC-FR), triple network-central executive (TN-CE), triple network-default mode (TN-DM), and triple network-salience (TN-SA). (D) Illustration of the spatiotemporal pattern detection algorithms used in this study: quasi-periodic patterns (QPPs, left), and complex principal component analysis (cPCA, right).

To distinguish the timing of scans relative to disease transition, we use the PRE and POST notation, where PRE refers to the scan before transition to a more advanced disease stage and POST refers to the scan after the transition has occurred. Within these groups, subjects in sNC, uNC, and sMCI initially do not exhibit clinical symptoms of dementia (DAT) during the study window. In contrast, pMCI and sDAT represent the DAT spectrum that spans from clinically asymptomatic to severely impaired. The choice of these groups was influenced by the availability of scan data, with Group 1 contributing hundreds of hours of fMRI data and Group 2 providing significantly fewer available hours (Table I). This framework allows us to systematically examine disease related changes in functional connectivity across both stable and transitioning cohorts. For further longitudinal information on Group 2, see Fig. 2. For information on preprocessing, quality assurance, and group level spatially constrained independent component analysis (scICA) using the GIFT toolbox, see [10] and Fig. 1A.

**Fig. 2:**
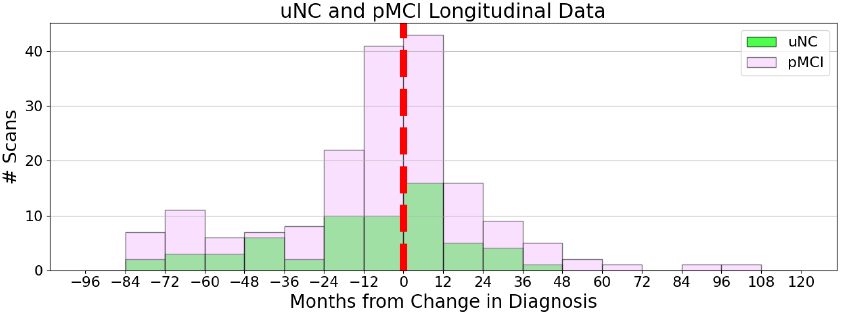
Longitudinal Demographics of Group 2. Histograms of available scans from longitudinal subjects who changed diagnosis during the duration of the research project. (Green) Transition from NC to MCI: uNC, (Purple) Transition from MCI to DAT: pMCI. The red line indicates the “Month 0” or the transition point where that session changed the diagnosis.

**TABLE I:**
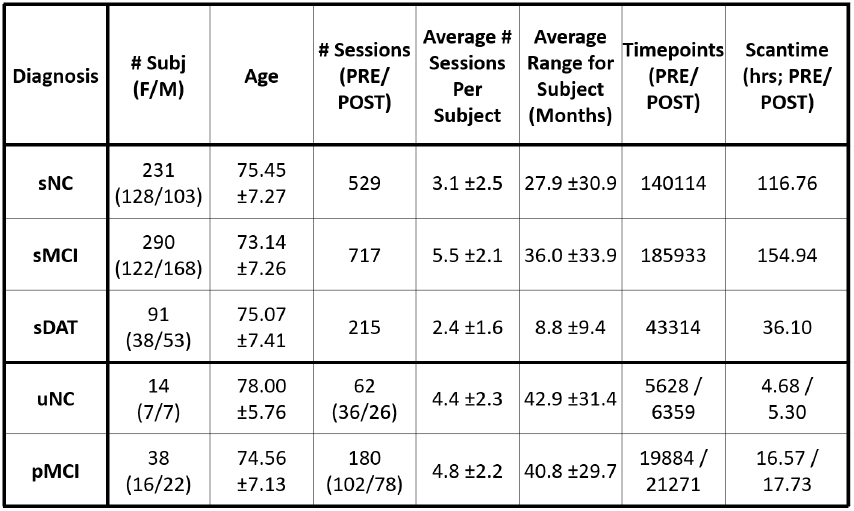
Demographics of the data across the two research Groups over longitudinal progression.

| Diagnosis | # Subj (F/M) | Age | # Sessions (PRE/ POST) | Average # Sessions Per Subject | Average Range for Subject (Months) | Timepoints (PRE/ POST) | Scantime (hrs; PRE/ POST) |
| --- | --- | --- | --- | --- | --- | --- | --- |
| sNC | 231 (128/103) | 75.45 ± 7.27 | 529 | 3.1 ± 2.5 | 27.9 ± 30.9 | 140114 | 116.76 |
| sMCI | 290 (122/168) | 73.14 ± 7.26 | 717 | 5.5 ± 2.1 | 36.0 ± 33.9 | 185933 | 154.94 |
| sDAT | 91 (38/53) | 75.07 ± 7.41 | 215 | 2.4 ± 1.6 | 8.8 ± 9.4 | 43314 | 36.10 |
| uNC | 14 (7/7) | 78.00 ± 5.76 | 62 (36/26) | 4.4 ± 2.3 | 42.9 ± 31.4 | 5628 / 6359 | 4.68 / 5.30 |
| pMCI | 38 (16/22) | 74.56 ± 7.13 | 180 (102/78) | 4.8 ± 2.2 | 40.8 ± 29.7 | 19884 / 21271 | 16.57 / 17.73 |

### B. NeuroMark Atlas

The NeuroMark v2.2 atlas [11] available in the GIFT software (http://trendscenter.org/software/gift) [12] and at http://trendscenter.org/data provides a replicable, data driven framework for analyzing functional connectivity in resting state fMRI, organizing functional brain networks into seven primary domains and fourteen subdomains (Fig. 1C). Each domain encapsulates ICNs, reflecting specialized brain roles in sensory processing, cognition, and emotion.

### C. QPP and cPCA Algorithms

We quantified large scale spatiotemporal dynamics using two complementary approaches: quasi periodic patterns (QPPs) and complex principal component analysis (cPCA). QPPs were detected using the QPPLab software package [13], which is designed to identify recurrent spatiotemporal patterns in blood oxygenation level dependent (BOLD) fMRI signals. Following prior work [14], the window length for QPP detection was set to 24 seconds (Fig. 1D), corresponding to approximately one cycle of infraslow oscillatory activity (0.01 to 0.15 Hz) and ensuring that a full cycle of the recurrent pattern is captured. QPPLab implements a seed-free, data driven sliding template algorithm that iteratively refines a spatiotemporal template by correlating it with the preprocessed ICN time courses and updating the template based on segments with the highest correlation. This procedure identifies repeating, phase coupled sequences of activity without assuming static connectivity across the scan. Compared to static functional connectivity or generic dynamic functional connectivity approaches, QPP analysis provides a compact representation of robust, recurrent patterns that have been shown to be reproducible across individuals, conditions, and species [14]–[17] and sensitive to disease related network alterations [7], [12], [18]. Consistent with prior work, we focused on the dominant QPP following global signal regression, which typically reflects alternating BOLD fMRI modulations between task positive and default mode networks [19]–[21]

To obtain a complementary pattern-based representation that captures both amplitude and phase relationships, we applied cPCA to the same global signal regression ICN time courses [7]. For each ICN *β*, an analytic signal *Z*_*β*_(*t*) = *I*(*β, t*) + *i* ℋ (*I*(*β, t*)) was constructed via the Hilbert transform, and a complex correlation matrix **R** was formed with entries 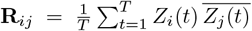, where 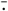 denotes complex conjugation. Eigen decomposition of **R** yielded complex spatial loadings |**u**_*k*_| and component time courses 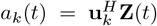. We focused on the leading component following global signal regression, whose magnitude **u**_1_ and phase arg(**u**_1_) define amplitude and phase delay maps across ICNs and capture a dominant phase coherent spatiotemporal pattern closely related to QPPs [7]. Further methodological details and applications of this approach to resting state fMRI can be found in [7].

### D. Network Integrity and Functional Connectivity Differences

The network level analysis begins with a set of cohort specific QPP templates *Q*_*i*_, where each *Q*_*i*_ ∈ ℝ^*N ×T*^ represents the dominant QPP for cohort *i, N* is the number of ICNs, and *T* is the template window length. For each cohort, we compute an ICN by ICN correlation matrix *C*_*ii*_ = corr(*Q*_*i*_, *Q*_*i*_), and cross cohort similarity matrices *C*_*ij*_ = corr(*Q*_*i*_, *Q*_*j*_), *i j*, where correlations are computed for each ICN pair by correlating their time courses within the QPP window (Fig. 3C). Deviations in QPP based network integrity relative to a reference cohort are summarized by Δ*C*_*ij*_ = *C*_*ii*_ − *C*_*ij*_, providing a matrix valued measure of how strongly QPP related functional connectivity in cohort *j* departs from the reference (Fig. 3D).

**Fig. 3:**
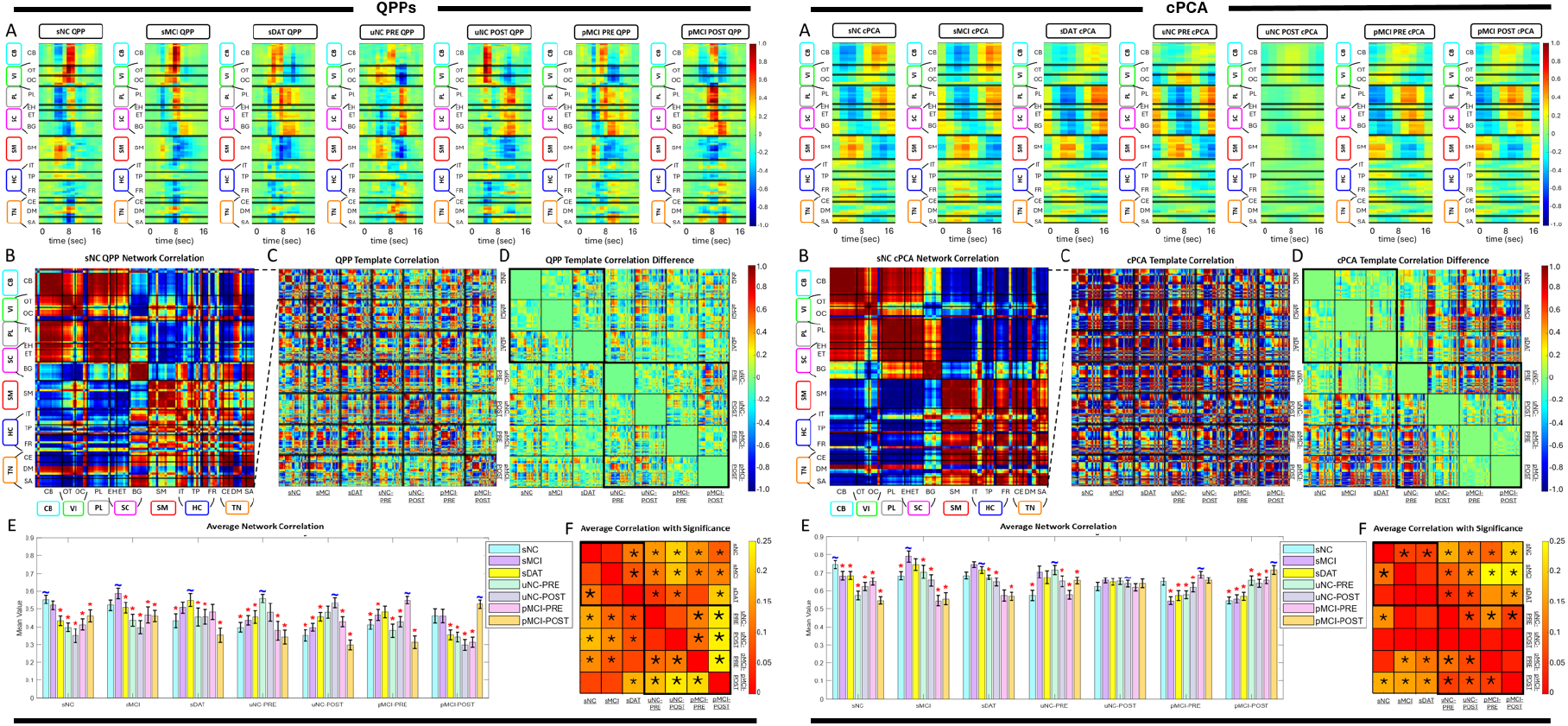
Evaluation of QPP and cPCA templates. (Left Panels A–F) illustrate the QPP based analysis, and (Right Panels A–F) illustrate the corresponding cPCA based analysis. In each set, (A) shows the group specific template for all ADNI diagnosis groups, with networks (rows) ordered by NeuroMark domains. (B) shows the network correlation matrix for the sNC template. (C) shows correlations between templates across disease progression. (D) uses the self correlation of each template as a baseline to display differences in network connectivity between groups, with bold black borders delineating Group 1 and Group 2 blocks. (E) summarizes the average correlation across network dynamics between templates, with red asterisks indicating significant differences in template correlations. (F) shows the results of Kruskal Wallis tests assessing differences in functional connectivity between ICNs across template correlations, where an asterisk marks network pairs with p-values less than *α* = 0.05.

In parallel, we derive cPCA based metrics by taking the magnitude of the leading complex spatial loading vector **u**_*i*_ ∈ ℂ^*N*^ for each cohort, yielding ICN level engagement scores **l**_*i*_ = |**u**_*i*_|. These magnitudes are then averaged within intrinsic connectivity domains to obtain domain level measures *L*_*i*_(*d*), which serve as cPCA based engagement indices for each functional domain. Statistical significance for both QPP and cPCA derived measures is assessed using the Kruskal Wallis test, which does not assume normality and is well suited to functional connectivity and component loading data [22]. The Kruskal–Wallis tests were applied to the distributions of ICN–ICN correlation values within each NeuroMark domain, treating each ICN pair as an observation. All statistical inferences are therefore made at the network-level rather than at the subject-level. We use *α* = 0.05 and, when the Kruskal Wallis test indicates a significant effect of group, apply Dunn’s post hoc test with Bonferroni correction for multiple comparisons to identify pairwise group differences (Fig. 3E,F). This framework enables robust comparisons of QPP based network integrity and cPCA derived network engagement across AD stages while accommodating non-Gaussian distributions and unequal variances.

### E. Data and Code Availability

The dataset is publicly available via the ADNI database (https://adni.loni.usc.edu/). Data processing was conducted in MATLAB, with visualization performed in MATLAB and Python. The QPP templates, analysis code, and brain spatial overlay videos can be accessed at https://github.com/GT-EmoryMINDlab/ADNI_Spatiotemporal_Templates.

## III. RESULTS

### A. QPP Derived Network Alterations Across AD Progression

QPP templates showed a systematic degradation of network integrity across both stable (Group 1: sNC, sMCI, sDAT) and transitioning (Group 2: uNC, pMCI) cohorts (Fig. 3, Fig. 4, Fig. 5). In Group 1, early changes from sNC to sMCI were largely confined to the paralimbic (PL), subcortical extended thalamic (SC-ET), and higher cognition (HC) domains, with prominent effects in frontal and insular-temporal networks. As subjects progressed to sDAT, QPP derived self-correlations revealed additional disruption in cerebellar (CB), visual (VI), and temporoparietal-HC networks, along with growing impairment in the triple network (TN; default and central executive subnet domains), indicating a shift from primarily limbic frontal alterations to more widespread breakdown of sensory, cerebellar, and large-scale cognitive control systems. Subcortical involvement remained a distinctive feature of the QPP results, with extended thalamic and hippocampal networks showing robust deterioration not captured by static functional connectivity alone.

**Fig. 4:**
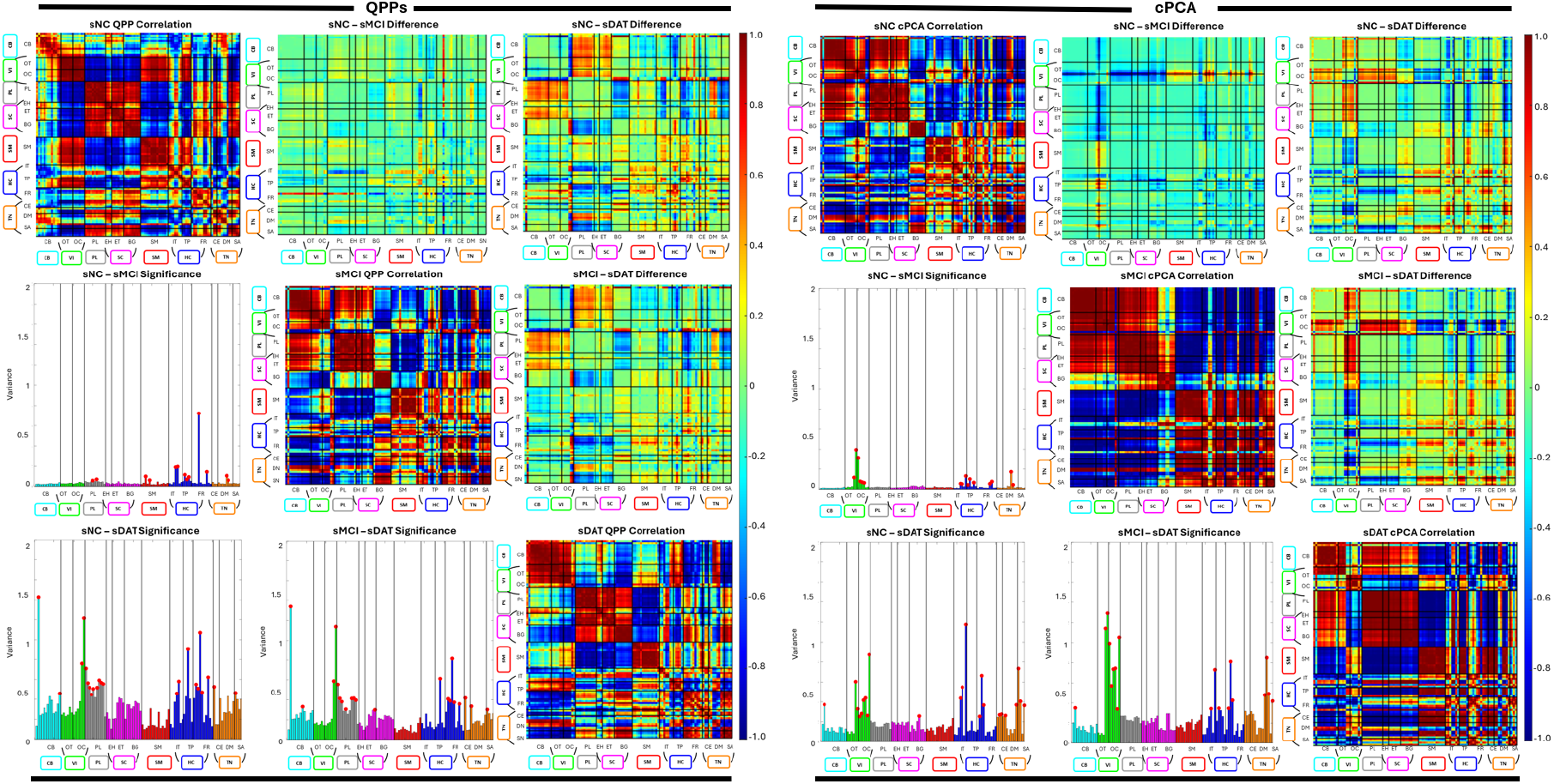
Evaluation of QPP and cPCA templates for Group 1. Left panels show results for QPP templates and right panels show results for cPCA templates. For each cohort, the matrices summarize network correlation differences with networks ordered by NeuroMark domains. The diagonal entries give within network correlations for the cohort specific template. The upper triangle shows differences in correlations between templates, and the lower triangle shows variance across networks, with red dots indicating network pairs with significant differences.

**Fig. 5:**
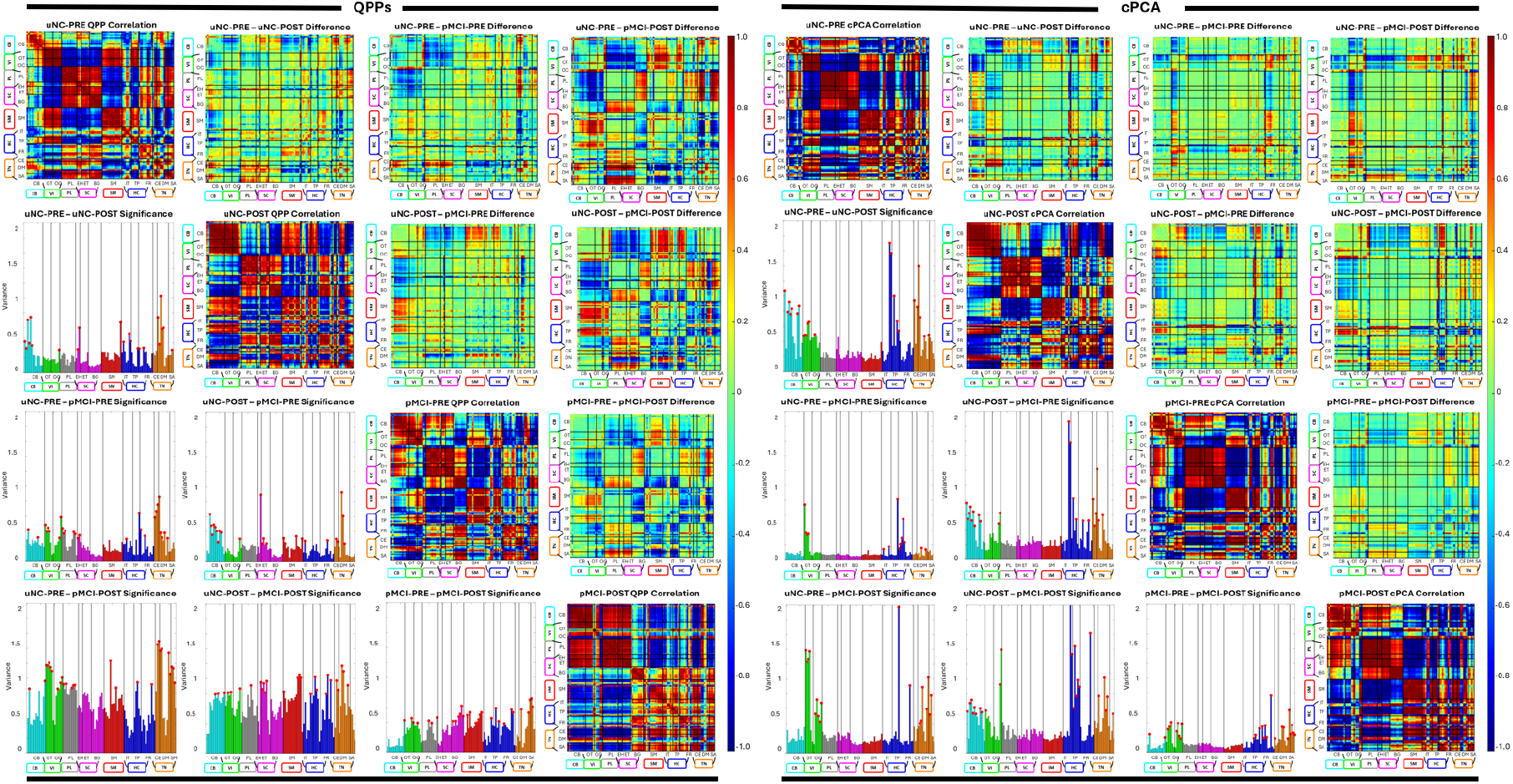
Evaluation of QPP and cPCA templates for Group 2. Left panels show results for QPP templates and right panels show results for cPCA templates in the transitioning cohorts. For each cohort, the matrices summarize network correlation differences with networks ordered by NeuroMark domains. The diagonal entries give within network correlations for the cohort specific template. The upper triangle shows differences in correlations between templates, and the lower triangle shows variance across networks, with red dots indicating network pairs with significant differences.

In Group 2, QPP analysis indicated that similar patterns emerged earlier and more abruptly in transitioning cohorts (Fig. 4). uNC subjects who later converted to MCI already showed QPP disruptions in cerebellar and visual domains, alongside alterations in higher cognition and triple network regions. With progression into pMCI, these changes expanded to include stronger paralimbic, subcortical extended hippocampal and thalamic (SC-EH), and sensorimotor (SM) involvement. Across uNC PRE, uNC POST, pMCI PRE, and pMCI POST, the burden of QPP based network disruption increased monotonically, with sensory (VI, SM), deep subcortical (SC), and triple network systems showing the clearest preclinical and progressive changes (Tables II–II Together, the QPP findings delineate an AD trajectory in which early limbic frontal and cerebellar visual disruptions emerge prior to diagnosis, followed by progressive breakdown of sensorimotor, subcortical, and large-scale control networks.

**TABLE II:**
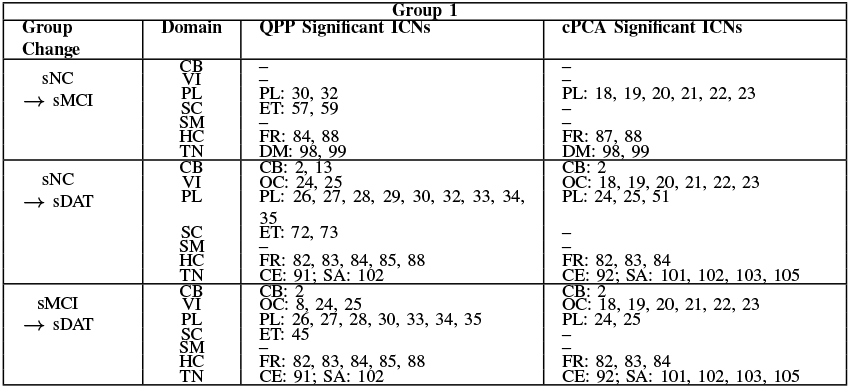
Significant ICNs Across Group 1.

### B. cPCA Derived Spatiotemporal Patterns

cPCA revealed complementary spatiotemporal alterations in large scale patterns, quantified using the domain averaged magnitudes *L*_*i*_(*d*) defined in the network integrity and functional connectivity differences subsection (Fig. 3, Fig. 4, Fig. 6). In Group 1, the leading cPCA pattern in sNC emphasized coherent structure spanning default mode, temporoparietal, and higher order cognitive networks, with anti-phase relationships to task-positive and sensorimotor systems. From sNC to sMCI, cPCA identified early changes concentrated in the paralimbic, higher cognition, and triple network domains, consistent with the QPP results but with fewer detectable effects in cerebellar and visual systems at this stage. With progression to sDAT, cPCA showed clear degradation of this pattern: amplitude decreased in default mode and frontal regions, visual involvement became more pronounced, and cerebellar contributions emerged, whereas subcortical alterations remained less prominent than in the QPP analysis. Overall, cPCA emphasized a gradual loss of spatially organized, phase coherent dynamics within higher order and visual networks, with later cerebellar involvement but minimal subcortical signal in stable cohorts.

**Fig. 6:**
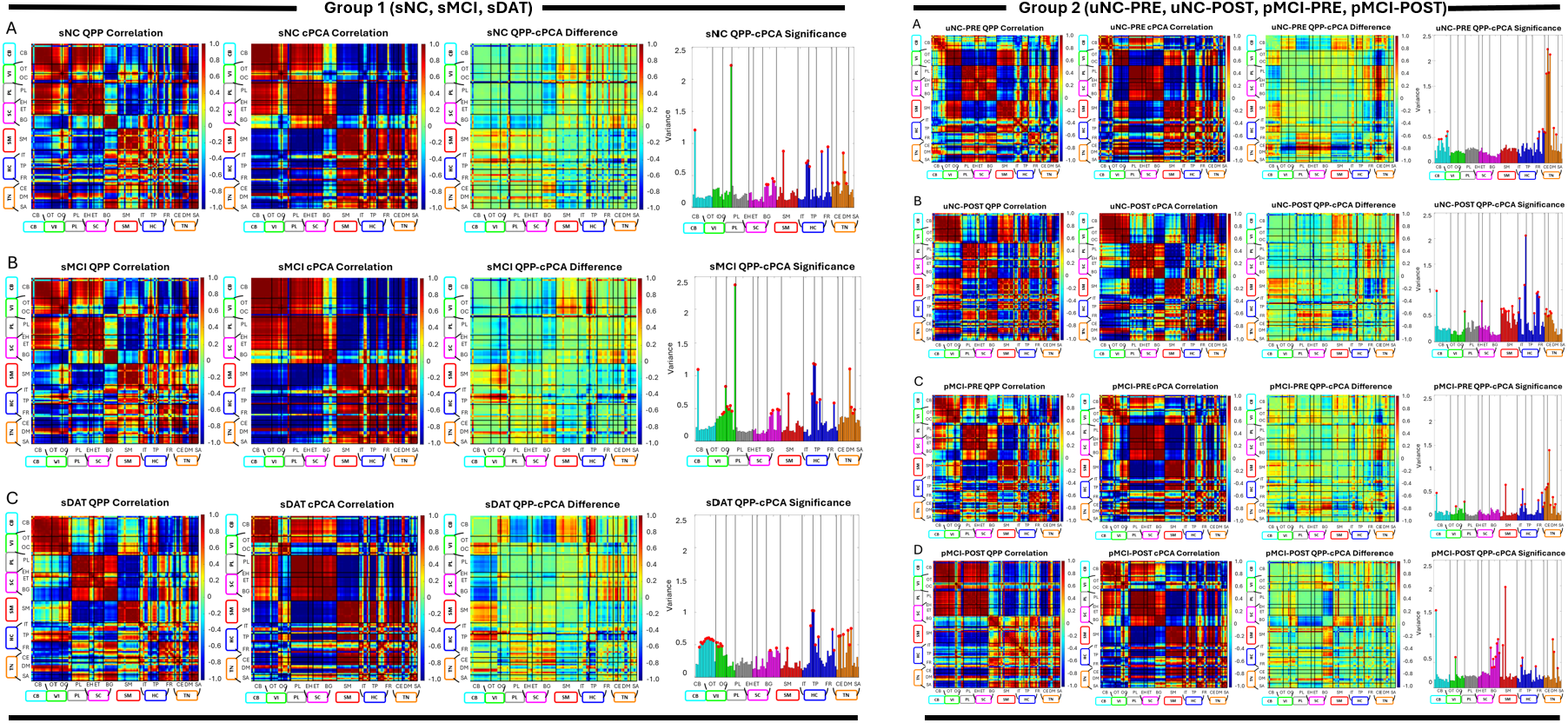
Comparison of QPP and cPCA for Groups 1 and 2. Left panels show stable cohorts in Group 1 (rows: sNC, sMCI, sDAT) and right panels show transitioning cohorts in Group 2 (rows: uNC PRE, uNC POST, pMCI PRE, pMCI POST). For each row, the four columns depict, from left to right: (1) QPP derived correlation, (2) cPCA derived correlation, (3) the difference between QPP and cPCA results, and (4) networks with statistically significant differences between the two methods. Together, these comparisons highlight where QPP and cPCA diverge in their ability to capture changes in functional connectivity across disease progression.

In Group 2, cPCA results were broadly consistent with the QPP based trajectory but highlighted some method specific differences. Transitioning subjects showed early alterations in cerebellar and visual domains, as well as in higher cognition and triple network components, echoing the QPP findings. However, paralimbic changes were weaker or absent in several transitions, and subcortical dysfunction emerged later and less consistently than in the QPP results. Across uNC PRE, uNC POST, pMCI PRE, and pMCI POST, cPCA loading magnitude decreased in frontal and temporoparietal networks and increased in variability within visual and cerebellar systems, indicating a progressive loss of coherent large scale propagation.

### C. Relationship Between QPP and cPCA Metrics

Comparing QPP and cPCA derived measures revealed both convergent and method specific patterns (Fig. 6, Tables II–III). At the domain level, both approaches consistently identified early and sustained involvement of higher cognition and triple network systems across AD stages, as well as progressive degradation in visual and cerebellar networks in later disease. Domains showing the largest QPP based declines in network integrity (for example PL, HC, TN, and later VI and CB) tended to exhibit reduced cPCA loading magnitude and less coherent spatial organization, such that preserved QPP structure generally coincided with stronger, more organized spatial patterns.

**TABLE III:**
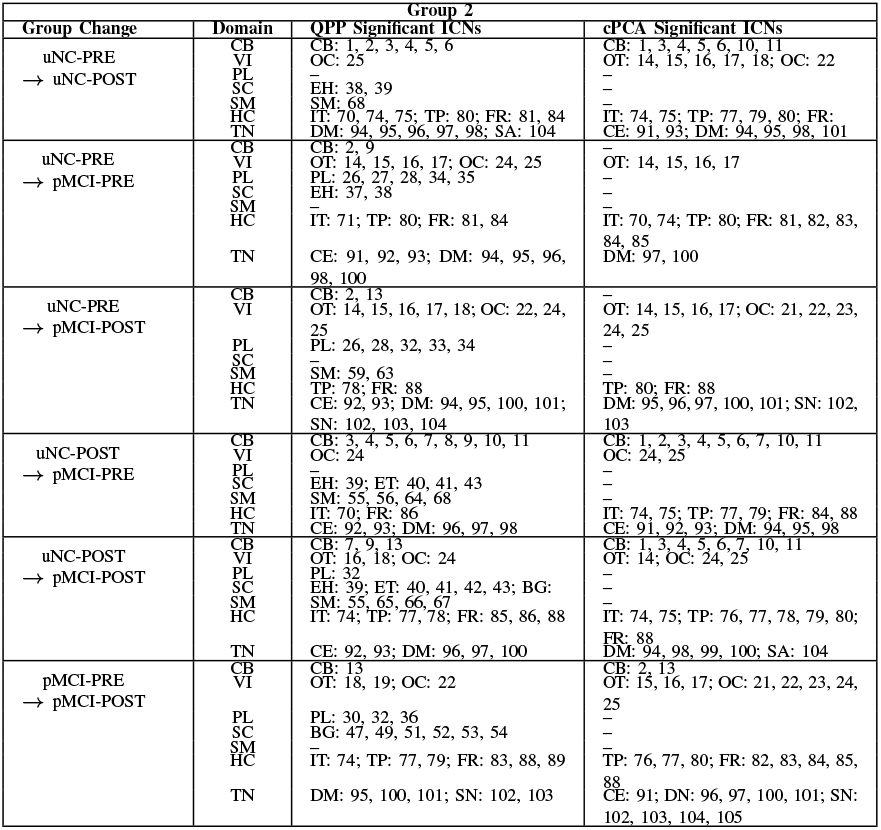
Significant ICNs Across Group 2.

Differences between the methods were most apparent in paralimbic and subcortical domains. QPPs detected robust and early limbic and subcortical alterations in both stable and transitioning cohorts, while cPCA was relatively insensitive to subcortical changes and captured fewer PL effects, particularly in Group 2. Conversely, cPCA often highlighted more distributed and persistent visual disruptions than QPPs, especially within occipital and occipitotemporal networks.

In transitioning cohorts, networks that showed early QPP changes (CB, VI, and TN) also displayed early cPCA deviations, with pMCI POST patterns approaching those of sDAT. These cross method comparisons indicate that QPP and cPCA metrics co vary in key domains yet also capture distinct aspects of domain vulnerability across the AD spectrum.

## IV. DISCUSSION

### A. QPP and cPCA Based Views of Network Degeneration

This study leveraged QPPs and cPCA to characterize spatiotemporal network alterations across the Alzheimer’s disease spectrum, extending our prior QPP only analysis [6]. Both approaches converged on a progression in which large scale networks become progressively less coordinated as disease severity increases, while highlighting partially distinct aspects of that degeneration.

In the stable cohorts, QPP derived measures revealed a sequence of domain involvement beginning with paralimbic, subcortical, and higher cognition systems and expanding to include cerebellar, visual, sensorimotor, and triple network domains in sDAT. cPCA identified a parallel degradation of a dominant phase coherent pattern spanning Default, temporoparietal, and higher order cognitive networks, with later involvement of visual and cerebellar systems. In the transitioning cohorts, both methods detected early deviations in cerebellar, visual, higher cognition, and triple network domains prior to formal diagnostic conversion, followed by further deterioration after conversion. Together, these results suggest that AD is characterized not only by reduced static connectivity but also by a loss of organized, recurrent spatiotemporal patterns and coherent propagation across distributed networks.

### B. Stable Versus Transitioning Cohorts and Early Network Vulnerability

Comparing stable (Group 1) and transitioning (Group 2) cohorts provides additional insight into the timing of these changes. The stable groups followed an expected progression from relatively preserved dynamics in sNC to widespread disruption in sDAT. By contrast, uNC and pMCI subjects already showed QPP based alterations in cerebellar, visual, higher cognition, and triple network domains before formal diagnostic conversion, with further deterioration after conversion. cPCA revealed similar early deviations in the same domains, although subcortical and paralimbic effects were generally weaker than in the QPP analysis.

These findings support a sequence in which limbic, subcortical, and higher cognition networks are affected early, followed by increasingly prominent visual, cerebellar, and sensorimotor abnormalities. The fact that both QPP and cPCA metrics show detectable deviations in uNC and pMCI strongly suggests that spatiotemporal network signatures carry preclinical information beyond conventional static connectivity, reinforcing the potential of these approaches for early detection and risk stratification. In a future clinical pipeline, such metrics could be incorporated as predictors in longitudinal mixed-effects or survival models to estimate individual conversion risk.

### C. Complementary Roles of QPPs and cPCA

Although QPP and cPCA converge on many of the same domains, their differences are informative. QPPs were more sensitive to limbic and subcortical disruptions, capturing early degradation in paralimbic and extended thalamic and hippocampal systems that were less prominent in the cPCA patterns. This likely reflects the fact that QPPs are driven by recurrent, high-amplitude events and can highlight transient but functionally meaningful excursions in limbic and subcortical circuits that are embedded within, rather than cleanly separated by, variance-based decompositions. In contrast, cPCA emphasized higher order cortical and visual systems and provided a compact description of how a leading phase coherent pattern becomes weaker and less organized with disease progression [7]. Thus, QPPs provide an event-based, primarily amplitude-weighted view of recurrent patterns, whereas cPCA offers a variance-based, phase-sensitive description of the same large-scale modes.

Taken together, these results suggest a useful division of labor. QPPs characterize the presence and integrity of recurrent spatiotemporal patterns, whereas cPCA characterizes trait like axes of dysfunction in the underlying covariance and phase delay structure. The strong network level agreement between the two methods in key domains (higher cognition, triple network, visual, cerebellar), combined with their differing sensitivity to limbic and subcortical changes, argues for their use as complementary tools rather than competitors.

### D. Limitations and Future Directions

Several limitations warrant consideration. First, the temporal resolution of the ADNI fMRI data (TR = 3 s) constrains sensitivity to faster dynamics and may underestimate the richness of QPP and cPCA patterns. Second, the sample size of transitioning cohorts is modest, limiting power to detect more subtle subgroup differences and increasing uncertainty around individual trajectories. Third, both QPP and cPCA depend on preprocessing choices (for example global signal regression and ICN parcellation), which may bias the relative visibility of cortical and subcortical effects. Replication in higher resolution datasets, with alternative preprocessing strategies and parcellations, will be important for establishing robustness.

Future work should extend these analyses in several directions. Methodologically, combining QPP and cPCA with other data driven approaches such as coactivation patterns (CAPs) [23], [24], spatiotemporal autoencoders [25], or related deep models may yield richer, low dimensional representations of disease related dynamics. Clinically, applying this framework to other dementias, i.e., frontotemporal, Lewy body, and vascular dementias, could reveal distinct spatiotemporal signatures with differential diagnostic value. Finally, integrating QPP and cPCA derived metrics with structural MRI, PET, and electrophysiological measures could clarify how functional spatiotemporal disruption relates to amyloid and tau burden, atrophy, and altered electrophysiological rhythms, and whether the observed network sequence mirrors known patterns of protein spread and network disconnection in AD, moving toward multimodal biomarkers that better capture the complexity of AD progression.

## V. CONCLUSION

This study shows that QPPs and cPCA capture a coherent trajectory of spatiotemporal network disruption across the Alzheimer’s disease spectrum. Using resting state fMRI from stable (sNC, sMCI, sDAT) and transitioning (uNC, pMCI) cohorts, we find that limbic, subcortical, and higher cognition networks are affected early, with progressive involvement of visual, cerebellar, sensorimotor, and triple network systems as disease severity increases. Importantly, many of these alterations are detectable in transitioning subjects prior to formal diagnostic conversion, indicating that spatiotemporal network signatures carry preclinical information.

QPP derived metrics emphasize the breakdown of recurrent, phase coupled patterns and highlight robust limbic and subcortical involvement, while cPCA derived patterns provide a complementary component-based view of how dominant phase coherent patterns in higher order, visual, and cerebellar networks weaken and become less organized. The convergence of these approaches on key vulnerable domains, together with their differing sensitivity to specific systems, supports their use as complementary tools rather than alternatives. As such, QPP and cPCA based measures offer a dual method framework for functional biomarkers of AD progression, with potential applications in early detection, risk stratification, and future treatment monitoring.

## Notes

* Research reported in this publication was supported by Goizueta Alzheimer’s Disease Research Center and NIH 1R01AG062581.

### Competing Interest Statement

The authors have declared no competing interest.

